# Local Mitochondrial Physiology Defined by mtDNA Quality Guides Purifying Selection

**DOI:** 10.1101/2025.08.08.669275

**Authors:** Felix Thoma, Romina Rathberger, Francesco Padovani, David Hörl, Kurt M. Schmoller, Christof Osman

## Abstract

The mitochondrial genome (mtDNA) encodes essential subunits of the electron transport chain and ATP synthase. Mutations in these genes impair oxidative phosphorylation, compromise mitochondrial ATP production and cellular energy supply, and can cause mitochondrial diseases. These consequences highlight the importance of mtDNA quality control (mtDNA-QC), the process by which cells selectively maintain intact mtDNA to preserve respiratory function.

Here, we developed a high-throughput flow cytometry assay for *Saccharomyces cerevisiae* to track mtDNA segregation in cell populations derived from heteroplasmic zygotes, in which wild-type (WT) mtDNA is fluorescently labeled and mutant mtDNA remains unlabeled. Using this approach, we observe purifying selection against mtDNA lacking subunits of complex III (*COB*), complex IV (*COX2*) or the ATP synthase (*ATP6*), under fermentative conditions that do not require respiratory activity. By integrating cytometric data with growth assays and simulations, we show that the decline of Δ*atp6* mtDNA reflects both intracellular selection for intact genomes and a proliferative disadvantage of cells lacking ATP synthase. In contrast, loss of Δ*cob* and Δ*cox2* mtDNA is primarily driven by selective maintenance of functional genomes.

In heteroplasmic cells containing both intact and mutant mtDNA, fluorescent reporters revealed local reductions in ATP levels and membrane potential (ΔΨ) near mutant genomes, indicating spatial heterogeneity in mitochondrial physiology that reflects local mtDNA quality. Disruption of the respiratory chain by deletion of nuclear-encoded subunits (*RIP1, COX4*) abolished these physiological gradients and impaired mtDNA-QC, suggesting that local bioenergetic differences are required for selective recognition.

Together, our findings support a model in which yeast cells assess local respiratory function as a proxy for mtDNA integrity, enabling intracellular selection for functional mitochondrial genomes.

## Introduction

Mitochondria contain their own genome, mitochondrial DNA (mtDNA), which is present in multiple copies in each cell. mtDNA encodes a limited but critical subset of mitochondrial proteins, which are integral subunits of the respiratory chain. Mutations in these genes often result in respiratory dysfunction, contributing to the pathogenesis of a wide range of diseases [1–3]. Purifying selection, which has been shown to act in the germline of mammalian cells and *Drosophila melanogaster*, acts to preserve mitochondrial DNA (mtDNA) integrity, by preventing maintenance of mutant mtDNA copies [4–9].

Our previous work demonstrated that even the single-celled yeast *Saccharomyces cerevisiae* can distinguish between intact and mutant mtDNA, establishing it as a powerful model for studying mitochondrial DNA quality control (mtDNA-QC) [10]. Remarkably, this quality control operates within a fused, continuous mitochondrial network and does not require the fission protein Dnm1. We further showed that selection against mutant mtDNA depends on intact mitochondrial cristae, which limit the diffusion of mtDNA-encoded proteins and thus preserve a spatial link between an mtDNA molecule’s genotype and its encoded proteins. These findings support the “sphere of influence” model [10, 11], which proposes that cristae-mediated compartmentalization of the mitochondrial inner membrane enables the emergence of distinct local physiological states that reflect the underlying mtDNA quality. Such spatial heterogeneity may in turn enable selective recognition of defective genomes. However, whether mutant mtDNA can indeed generate localized physiological differences within a continuous mitochondrial network that also harbors intact genomes has remained a critical open question.

To date, our investigations have primarily focused on a competitive context in which WT mtDNA coexists with mutant mtDNA lacking the *COB* gene, which encodes a core subunit of complex III [10, 12]. However, it remains unclear whether mutations affecting subunits of complex IV or the ATP synthase are similarly subject to selective elimination. This question is relevant because defects in different components of the respiratory chain have distinct bioenergetic consequences. Loss of complexes III or IV compromises the electron transport chain’s ability to generate a proton gradient, requiring reverse operation of the ATP synthase to maintain the membrane potential (ΔΨ), a process that consumes ATP and likely leads to ATP depletion [13–15]. In contrast, loss of the mtDNA-encoded Atp6, a F_O_ subunit of the ATP synthase, does not directly impair proton pumping by complexes III and IV. Because protons can no longer re-enter the matrix via the ATP synthase in the absence of Atp6, dissipation of ΔΨ is impaired, potentially resulting in membrane hyperpolarization, while ATP synthesis is abolished [16, 17]. However, the absence of ATP synthase has been reported to cause assembly defects in complex IV, further compounding bioenergetic dysfunction [18, 19]. Taken together, these observations suggest that the physiological consequences of specific mtDNA mutations may differ, and how these differences influence purifying selection remains an open question.

In this study, we developed a rapid analysis pipeline, termed FAST (Flow cytometry Analysis for Segregation Tracking), which combines micromanipulation with flow cytometry to monitor the segregation of WT and mutant mtDNA variants over multiple generations. Using FAST, we analysed the competitive dynamics between intact mtDNA and mutant genomes lacking subunits of complex III, complex IV, or the ATP synthase in populations derived from heteroplasmic zygotes containing both genome types. We further investigated whether ΔΨ and ATP levels differ locally within the mitochondrial network depending on the underlying mtDNA genotype. Finally, we tested whether mtDNA quality control requires a functional respiratory chain by assessing whether global impairment of oxidative phosphorylation disrupts the selective retention of intact mtDNA.

## Results

### Development of a Rapid Pipeline for mtDNA Segregation Analysis

In the following, we describe the development of a novel flow cytometry–based pipeline that enables quantitative analysis of mtDNA variant segregation in cell populations derived from single heteroplasmic zygotes carrying two distinct mitochondrial genomes. This approach builds on previous microscopy-based strategies that employed heteroplasmic yeast zygotes containing mtDNA encoding the fluorescent fusion protein Atp6-mNeonGreen (mtDNA^*ATP6-NG*^) alongside a non-fluorescent mtDNA (“dark” mtDNA) [12].

First, we asked whether flow cytometry allows reliable discrimination between non-fluorescent cells and cells harboring mtDNA^*ATP6-NG*^. To this end, we analysed three distinct populations: (i) ‘dark’ cells containing mtDNA not encoding fluorescent proteins, (ii) mtDNA^*ATP6-NG*^ cells and (iii) a mixture of both (Fig 1A–C). Flow cytometric analysis revealed that mtDNA^*ATP6-NG*^ cells could be readily distinguished from dark cells. Although non-fluorescent and mtDNA^*ATP6-NG*^ cells exhibited similar cell size based on forward scatter (FSC-A), only the latter showed markedly increased fluorescence intensity. This difference was particularly evident in the mixed population, where the two cell types formed clearly separated clusters (Fig 1C).

**Fig 1.**
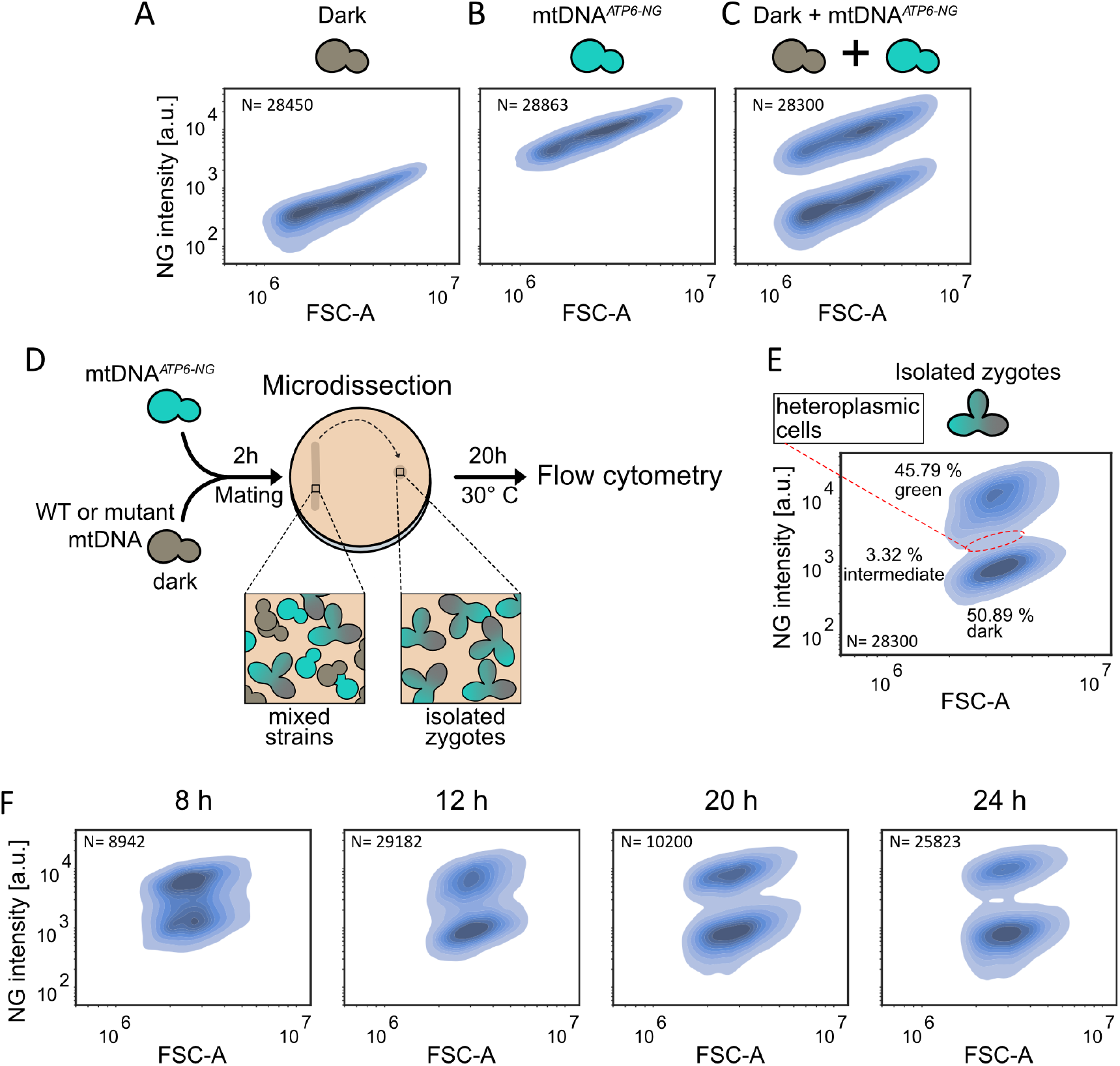
Flow cytometry based method to assess mtDNA quality control. Flow-cytometric density plot of yeast cells lacking NG **(A)**, containing mtDNA^*ATP6-NG*^ **(C)**, and a mixed population of both strains **(C)** with fluorescence intensity on the y-axis and forward scatter (FSC-A) on the x-axis. **(B)** Schematic of the FAST experimental workflow. **(E)** FAST density plot of mtDNA^*ATP6-NG*^ × mtDNA^ic^ heteroplasmic microcolonies after 20 h: the area highlighted with a red dashed circle indicates heteroplasmic cells; percentages of fluorescent, heteroplasmic, and non-fluorescent populations are annotated next to each cluster. **(F)** mtDNA^*ATP6-NG*^ cells were mated with mtDNA^ic^ cells. 310, 150, 50, and 40 zygotes were isolated to discrete coordinates on an agar plate and microcolonies were analysed 8, 12, 20, and 24 h post-mating, respectively.

In the next step, we tested whether flow cytometry enables rapid and quantitative analysis of mtDNA segregation from heteroplasmic founder cells. We crossed the mtDNA^*ATP6-NG*^ (intron-containing) strain with a WT strain harboring intact intron-containing mtDNA (mtDNA^ic^), which lacks a gene encoding a fluorescent protein, to generate heteroplasmic zygotes. Note that, depending on the experimental context, both intron-containing and intronless mtDNA (mtDNA^il^) variants are used throughout this study. After zygote formation on agar plates, we isolated 30 zygotes using a microdissection microscope and transferred them to a single separate area on the plate. The plates were then cultured for 20 hours to allow microcolony formation, after which the colony was collected and subjected to flow cytometry analysis (Fig 1D). Flow cytometry revealed clear mtDNA segregation, with 45.8% of cells displaying fluorescence and 50.9% showing no fluorescent signal (Fig 1E). Notably, the size distribution in Fig 1E is narrower than in the liquid-grown controls (Fig 1A–C), likely because the microcolonies came from cells grown on solid medium, while the controls were from log-phase liquid cultures, which show more size variation. While the segregation of cells into two clusters was evident, a subset of cells displayed intermediate fluorescence (3.32%, highlighted in Fig 1E). Those intermediate cells were defined based on a narrow fluorescence intensity window (±10% of the distance between peaks) centered around the local minimum between the two major peaks in the kernel density estimate (KDE) of the NG signal distribution (S1 FigB). We hypothesized that these might represent residual heteroplasmic cells. To explore transition from heteroplasmic to homoplasmic cells, we monitored the cell population over a time course at 8, 12, 20, and 24 hours after mating. Indeed, we observed a gradual shift toward two virtually completely separated discrete clusters: one fully fluorescent and one non-fluorescent (Fig 1F).

Our previous research had demonstrated that the first cell division of a zygote plays a major role in the segregation of mtDNA variants [10]. This led us to test if segregation might be faster when analysing populations derived from the first daughter cell of a heteroplasmic zygote, rather than populations derived from the zygote itself. To test this, we isolated the first daughter cells from 30 heteroplasmic zygotes, transferred them to a separate location on the agar plate, and examined mtDNA segregation after 20 hours (S1 FigD). This approach indeed revealed a more complete segregation of mtDNA variants compared to the analysis performed directly on the zygotes: the two clusters were more distinct, and the proportion of heteroplasmic cells fell from 3.32% to 1.8% (S1 FigA–C). Hence, we applied this procedure for all following mtDNA segregation experiments.

In summary, this workflow enables fast data acquisition and high-throughput, single-cell resolution analysis. We refer to this streamlined approach as FAST-Flow cytometry Analysis for Segregation Tracking.

### Competitive advantage of WT over mutant mtDNA

We next sought to determine whether mtDNA harboring deletions of different genes encoding subunits of different respiratory chain complexes would exhibit altered segregation patterns. In our previous studies, we demonstrated a pronounced preference for WT mtDNA inheritance when mating WT strains with strains harboring a deletion of the *COB* gene in their mtDNA (mtDNA^Δ*cob*^) using microscopic or genetic readouts [10, 12]. Since the mtDNA^Δ*cob*^ strain was derived from mtDNA lacking introns, we included mating experiments as controls in our FAST analysis in which strains containing mtDNA^*ATP6-NG*^ were crossed with strains containing either intact mtDNA^ic^ or mtDNA^il^ [20]. Using our FAST analysis, we confirmed the purifying selection against Δ*cob* mtDNA: mating WT mtDNA^*ATP6-NG*^ with mtDNA^Δ*cob*^ resulted in a significant skew towards cells containing intact mtDNA, with approximately 82.1% WT mtDNA^*ATP6-NG*^-containing cells and only around 17.9% dark cells (Fig 2B+D). In contrast, mating strains containing mtDNA^*ATP6-NG*^ with WT strains harboring intact mtDNA^ic^ or mtDNA^il^ resulted in a more balanced segregation (Fig 2A+D, S1 FigA). Notably, we observed a slight competitive advantage of mtDNA^ic^ over mtDNA^*ATP6-NG*^, resulting in a higher proportion of cells retaining the ‘dark’ mtDNA^ic^. This indicates that cells may counterselect against mtDNA^*ATP6-NG*^ to some extent. This finding suggests that Atp6-NG might be functionally compromised, despite the absence of detectable growth defects in strains harboring mtDNA^*ATP6-NG*^, which was assessed in previous studies [10].

**Fig 2.**
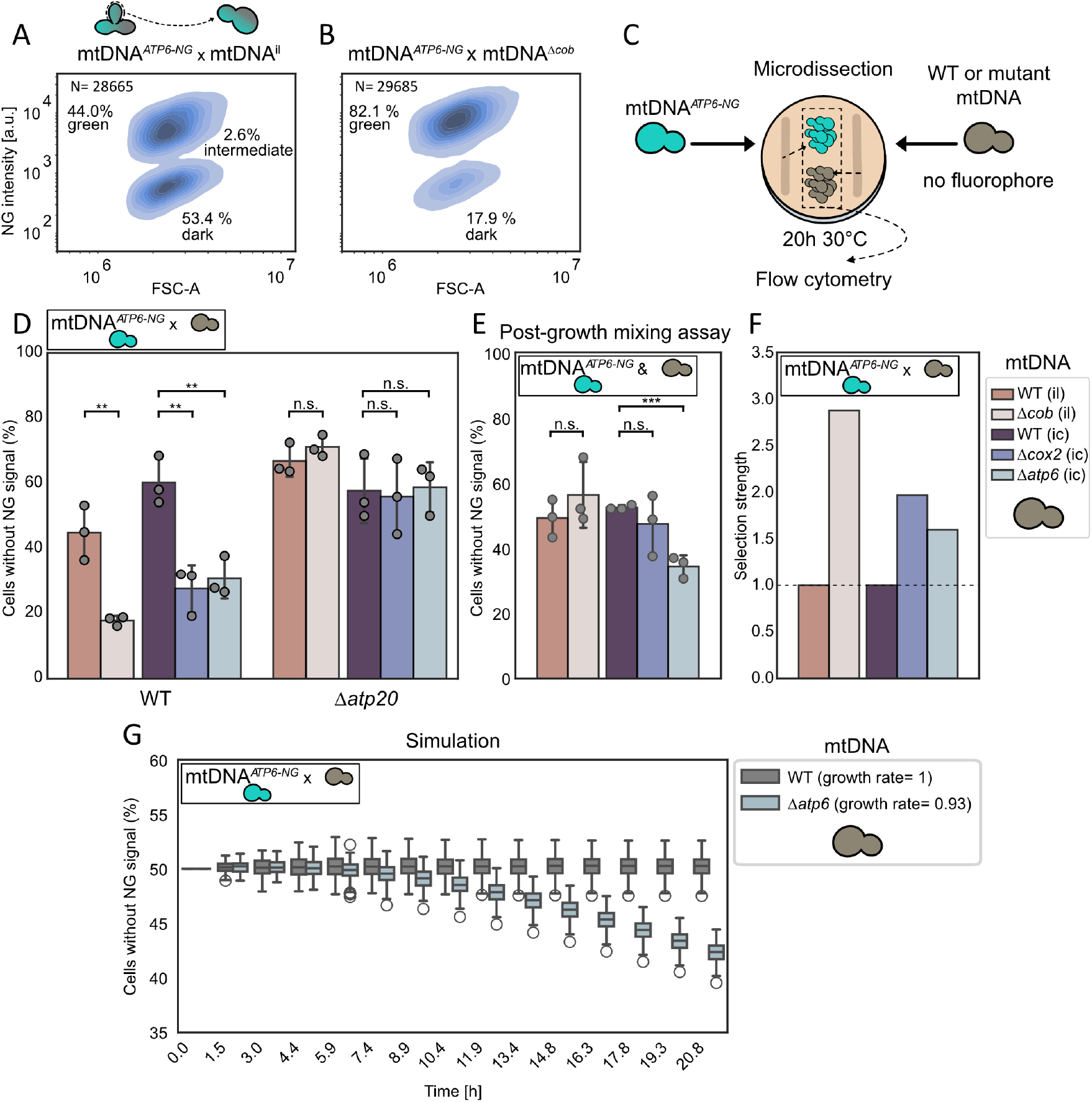
Competitive advantage of WT over mutant mtDNA. FAST results for matings of mtDNA^*ATP6-NG*^ with **(A)** mtDNA^il^ or **(B)** mtDNA^Δ*cob*^. 30 first daughter cells from formed zygotes were isolated by micromanipulation, cultured for 20 h, and analysed by flow cytometry. **(C)** Schematic of the post-growth mixing assay workflow: 30 cells of mtDNA^*ATP6-NG*^ or non-fluorescent strains were microdissected to a defined, separate position, cultured for 20 h, then pooled for flow-cytometric analysis. **(D)** Bar graph depicting the fraction of non-fluorescent cells for each FAST mtDNA^*ATP6-NG*^ mating combination in WT and Δ*atp20* strains. Individual replicates are indicated by dots. **(E)** Results of the post-growth mixing assay showing the proportion of non-fluorescent cells. **(F)** Calculation of selection strength derived from the data in panels (C) and (D) and then normalized to WT. **(G)** Simulation of mtDNA^*ATP6-NG*^ x mtDNA^Δ*atp6*^ mating without any mtDNA selection. Statistical significance in panels (D) and (E) was determined by unpaired Student’s t-test on the means of replicates (*P *<*0.05; **P *<*0.005; ***P *<* 0.001).

Next, we asked whether the observed selective disadvantage is specific to the *COB* gene or whether similar selection patterns apply to mutations in other mtDNA-encoded genes. To address this, we extended our analysis to mtDNA deletion strains lacking either *COX2* (mtDNA^Δ*cox2*^), which encodes a core subunit of complex IV, or *ATP6*, a core component of the ATP synthase complex (mtDNA^Δ*atp6*^) [21–23]. Like the mtDNA^Δ*cob*^ strain, mtDNA^Δ*cox2*^ and mtDNA^Δ*atp6*^ strains were unable to grow on non-fermentable carbon sources (S1 FigE). We then performed matings between strains containing intact mtDNA^*ATP6-NG*^ and strains harboring either mtDNA^Δ*cox2*^ or mtDNA^Δ*atp6*^. Both mtDNA^Δ*cox2*^ and mtDNA^Δ*atp6*^ contain introns, and thus results from the FAST analyses involving these strains must be compared to those from an mtDNA^*ATP6-NG*^ *×* mtDNA^ic^ cross. In both cases, the amount of non-fluorescent cells was reduced (27.6% and 30.8%, respectively), pointing to a preferential maintenance of intact mtDNA in the resulting colonies (Fig 2D).

We validated these findings by repeating the analysis using fluorescence microscopy instead of flow cytometry. For this analysis, individual colonies derived from the first daughter cells of heteroplasmic zygotes were imaged separately to assess cell-to-cell variability. This approach highlighted a stochastic element in the segregation process, with some colonies from heteroplasmic zygotes composed almost entirely of non-fluorescent (dark) cells and others containing predominantly fluorescent cells (S2 FigA+B). Nonetheless, when results were averaged across multiple colonies, they consistently converged on a reproducible mean, which was comparable to our findings obtained through FAST.

Strains carrying mutant mtDNA variants may exhibit slower growth rates, even on fermentable medium, compared to their corresponding WT strains, potentially contributing to a lower percentage of cells containing mutant mtDNA in populations resulting from heteroplasmic founder zygotes. For instance, the predominance of green-fluorescent cells after 20 hours might not necessarily indicate an intracellular selective advantage of the mtDNA^*ATP6-NG*^, but rather reflect a growth-rate advantage over their counterparts harboring mutant mtDNA.

Assessment of doubling times for strains containing either intact (mtDNA^il^ or mtDNA^ic^) or mutant (mtDNA^Δ*cob*^, mtDNA^Δ*cox2*^, or mtDNA^Δ*atp6*^) mtDNA in liquid medium did not reveal significant growth differences (S2 FigC+D). However, the FAST analysis is performed on solid medium, where spatially heterogeneous growth conditions within colonies may impose additional metabolic challenges [24–26]. To better reflect these conditions, we sought to assess the growth behavior of the different yeast strains under similar settings, and therefore repeated the FAST analysis without the mating step. Specifically, thirty haploid budding cells were isolated and deposited at a defined position on the agar plate, giving rise to a microcolony derived from these founder cells. This procedure was performed for all strains harboring the various mtDNA variants. After 20 hours of growth, the resulting colonies were harvested and pooled in combinations mimicking those used in the mating-based FAST experiments. The pooled populations were then immediately analysed by flow cytometry. We refer to this assay as ‘post-growth mixing assay’(Fig 2C).

For mixtures of WT mtDNA^*ATP6-NG*^ and either mtDNA^Δ*cob*^ or mtDNA^Δ*cox2*^ cells, approximately 50% of the population exhibited fluorescence, consistent with similar growth rates between strains (Fig 2E). Thus, the observed reduction of non-fluorescent mutant cells in the FAST experiments involving mtDNA^Δ*cob*^ or mtDNA^Δ*cox2*^ cannot be explained by growth differences. In contrast, mixtures of independently grown mtDNA^*ATP6-NG*^ and mtDNA^Δ*atp6*^ cells displayed a lower proportion of mtDNA^Δ*atp6*^ cells in the post-growth mixing assay, indicating a growth disadvantage for this strain under plate-based conditions. This growth defect likely contributes to a decreased percentage of mtDNA^Δ*atp6*^ cells in FAST analysis of populations derived from heteroplasmic zygotes and may lead to an overestimation of intracellular selection against mutant mtDNA.

In our post-growth mixing assay, cells were homoplasmic for either mtDNA^*ATP6-NG*^ or mtDNA^Δ*atp6*^, which is an extreme scenario that would arise if complete segregation of mtDNA would occur during the first cell division. In contrast, mtDNA segregation starts from heteroplasmy and proceeds gradually [12]. Hence, the growth disadvantage due to mtDNA^Δ*atp6*^ is likely minor at first and increases as cells approach homoplasmy. To more accurately examine the relative contributions of mtDNA quality control and the growth disadvantage observed in mtDNA^Δ*atp6*^-containing cells, we turned to a previously established mtDNA segregation simulation framework [12]. This simulation models mtDNA segregation within a population, and accounts for key aspects of mitochondrial dynamics, including fusion and fission, and mtDNA partitioning during cell division (S2 FigF). We modified the model by introducing a growth penalty in heteroplasmic cells containing both mtDNA^Δ*atp6*^ and intact mtDNA, assuming a linear relationship between the growth defect and the percentage of mtDNA^Δ*atp6*^ copies within the cell. To parameterize the model, we first determined the doubling time of 1.48 hours of mtDNA^*ATP6-NG*^ cells on plates by seeding 10, 20, or 30 cells and measuring cell numbers after 20 hours using flow cytometry (S2 FigE). Together with the fluorescence ratios in the post-growth mixing assay, this allowed us to calculate the relative growth rate of mtDNA^Δ*atp6*^ cells compared to WT cells of 0.93, which we then used as a growth penalty in the simulation. Incorporating these values into our simulations allowed us to estimate the expected proportion of mtDNA^*ATP6-NG*^ in the population after about 20 hours of growth, corresponding to the conditions used in the FAST analysis.

In these simulations, which assume no intracellular competitive disadvantage of the mtDNA^Δ*atp6*^, but simply take into account growth defects associated with the presence of mtDNA^Δ*atp6*^, the resulting population was composed of approximately 42.9% mtDNA^Δ*atp6*^ cells and 57.1% mtDNA^*ATP6-NG*^ cells (Fig 2G). In contrast, our FAST analysis revealed only 31% mtDNA^Δ*atp6*^ cells. This discrepancy suggests that while the growth disadvantage associated with mtDNA^Δ*atp6*^ contributes to the reduced representation of this mtDNA variant in segregation analyses, it cannot fully explain the observed decline. Thus, our data support the occurrence of active selection against mtDNA^Δ*atp6*^ within the population.

To quantify the selection strength against the different mutant mtDNA variants, we proceeded as follows: For mtDNA^Δ*atp6*^, the outcome of the simulation analysis was divided by the corresponding FAST result to correct for effects related to growth rate. Since mtDNA^il^, mtDNA^ic^, mtDNA^Δ*cob*^ and mtDNA^Δ*cox2*^ cells did not exhibit any growth defects in our post-growth mixing assay, this correction step was omitted for these mtDNA variants, and directly divided by the FAST results. Subsequently, values from crosses involving mutant mtDNA were normalized to the respective WT cross (mtDNA^*ATP6-NG*^ *×* mtDNA^il^ for mtDNA^Δ*cob*^, and mtDNA^*ATP6-NG*^ *×* mtDNA^ic^ for mtDNA^Δ*cox2*^ and mtDNA^Δ*atp6*^), yielding what we refer to as selection strength. A value of 1 indicates no selective advantage for either mtDNA variant due to intracellular selection, whereas values greater than 1 indicate preferential inheritance of one mtDNA variant over the other (Fig 2F). This analysis revealed stronger selection against the mtDNA^Δ*cob*^ and mtDNA^Δ*cox2*^ variants, while selection against the mtDNA^Δ*atp6*^ variant appeared comparatively weaker.

As shown previously, mitochondrial ultrastructure plays an important role in maintaining mtDNA-QC by ensuring that proteins remain localized near the genome from which they originate [10]. It has been proposed that this spatial organization allows each mitochondrial genome to supply proteins specifically to the respiratory chain complexes within its immediate vicinity, thereby creating a localized sphere of influence [10, 11].

In our previous work, we demonstrated that deletion of *ATP20*, a gene required for proper cristae formation, impairs mtDNA-QC against mtDNA lacking the *COB* gene [10]. To confirm these findings, to test our system, and to examine whether cristae formation is also critical for selection against mtDNA^Δ*cox2*^ and mtDNA^Δ*atp6*^, we performed FAST with strains lacking *ATP20*. Indeed, mtDNA-QC was strongly reduced in Δ*atp20* cells and segregation shifted towards similar percentages of mtDNA^*ATP6-NG*^ cells and cells containing mtDNA^Δ*cob*^, mtDNA^Δ*cox2*^ or mtDNA^Δ*atp6*^, implying that normal cristae formation via assembly of the dimeric ATP synthase [27] is critical for mtDNA-QC(Fig 2D,S2 FigH). Interestingly, the growth phenotype in mtDNA^Δ*atp6*^ strains compared to mtDNA^*ATP6-NG*^ was rescued in the background of Δ*atp20*, explaining why percentages of dark cells were virtually identical between mtDNA^*ATP6-NG*^ *×* mtDNA^ic^ and mtDNA^*ATP6-NG*^ *×* mtDNA^Δ*atp6*^ crosses in a Δ*atp20* background (S2 FigG).

Thus, our analysis reveals that *S. cerevisiae* cells can discriminate between intact mtDNA and mtDNA lacking genes encoding components of complexes III, IV or ATP synthase thereby promoting the maintenance of intact mtDNA. Notably, selection against mtDNA^Δ*atp6*^ appeared less efficient than against mtDNA^Δ*cob*^ or mtDNA^Δ*cox2*^. This finding is reminiscent of previous studies in mammalian systems, which also report reduced selective pressure on *ATP6* compared to other mtDNA-encoded genes [9, 28].

### Local differences of ΔΨ and ATP levels within mitochondria of heteroplasmic zygotes

According to the sphere-of-influence hypothesis, each mitochondrial genome primarily supplies its immediate surroundings with gene products [10, 11]. Although mitochondria form a continuous network with content equilibration of soluble matrix proteins, mtDNA and their membrane-bound gene products remain spatially constrained and do not mix rapidly. Thus, mtDNA deletions could locally impair respiration, leading to reduced ATP production and lower ΔΨ in their vicinity. To test this hypothesis, we next asked whether mitochondrial physiology indeed differs between regions of the network that contain intact versus mutant mtDNA.

As a prerequisite to testing whether local physiological differences reflect mtDNA quality, we first used fluorescence microscopy to assess whether differences in ΔΨ and ATP levels are detectable at the single-cell level in strains containing exclusively WT or mutant mtDNA. Relative ATP levels were quantified using the ratiometric sensor QUEEN-2m, which reports ATP concentration based on shifts in its excitation spectrum upon ATP binding [29, 30]. ΔΨ was assessed using tetramethylrhodamine methyl ester (TMRM), a positively charged, lipophilic dye that accumulates in mitochondria in a ΔΨ-dependent manner [31, 32]. To correct for mitochondrial content, TMRM fluorescence was normalized to the NG signal which was fused to the mitochondrial targeting sequence of subunit 9 of the *Neurospora crassa* ATP synthase. This marker is known to localize to mitochondria in a manner largely independent of ΔΨ [33]. WT cells exhibited significantly higher mitochondrial ATP levels and normalized TMRM signal compared to mtDNA^Δ*cob*^ and mtDNA^Δ*cox2*^ strains (Fig 3A-D). In mitochondria harboring mutant mtDNA, both ATP levels and ΔΨ were markedly reduced. These reductions resembled those observed in cells treated with carbonyl cyanide 3-chlorophenylhydrazone (CCCP), a protonophore that dissipates ΔΨ, or 2-deoxy-D-glucose (2-DG), which inhibits glycolysis and limits pyruvate supply to the TCA cycle, thereby reducing cellular ATP levels to below 1% [30, 34]. These findings indicate that the mutant strains exhibit severely compromised mitochondrial function. As expected, mtDNA^Δ*atp6*^ cells also showed a marked reduction in mitochondrial ATP levels relative to WT. Notably, ΔΨ was also significantly reduced in mtDNA^Δ*atp6*^ cells, despite the presence of all genes encoding the structural components necessary for the assembly of complexes III and IV, which in principle should be capable of generating a ΔΨ. This finding aligns with previous reports demonstrating that the absence of an assembled ATP synthase can impair complex IV stability, thereby compromising the respiratory chain’s ability to maintain ΔΨ [18, 19].

**Fig 3.**
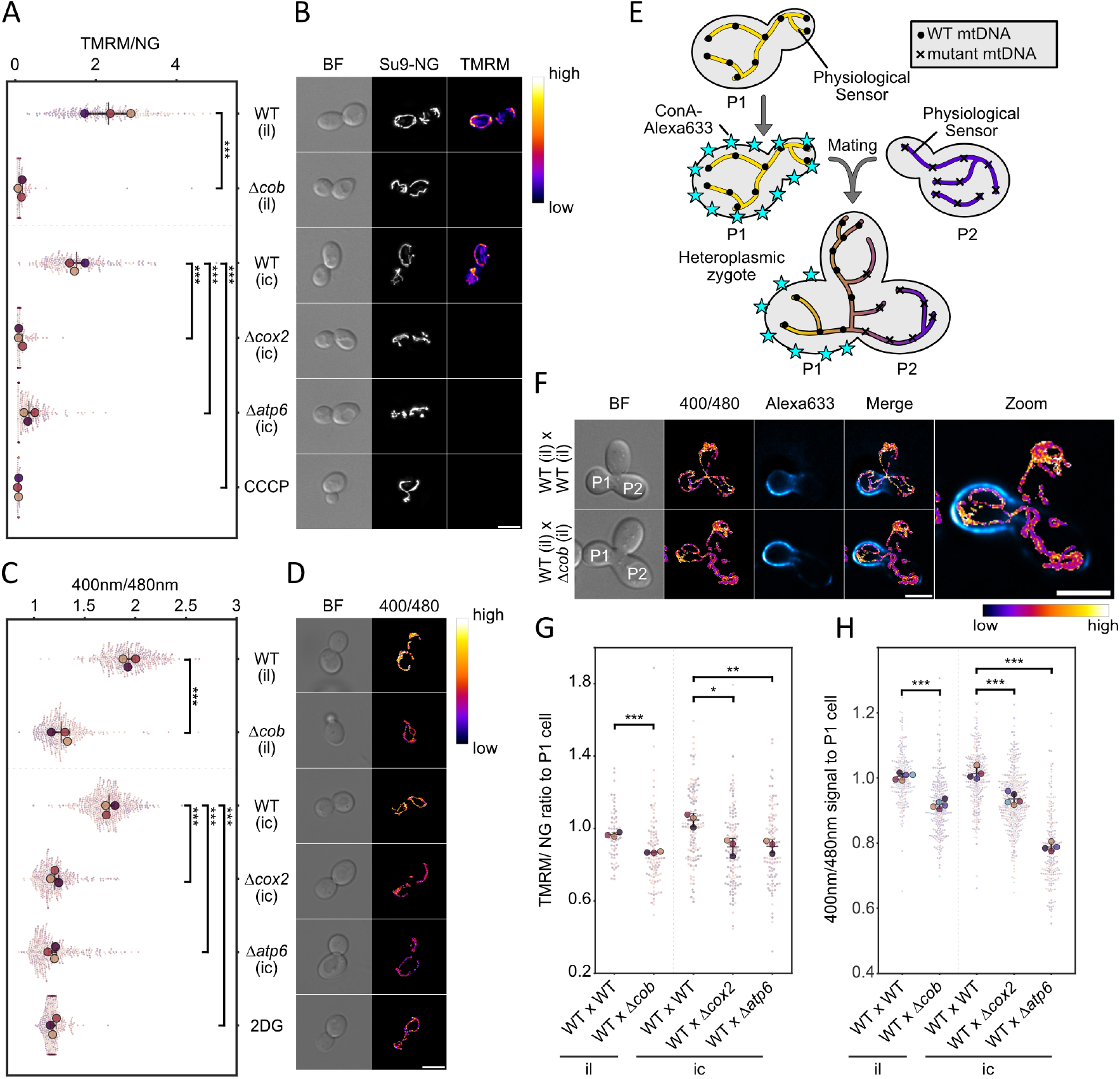
Local differences of ΔΨ and ATP levels within mitochondria of heteroplasmic zygotes. **(A)** Superplot of membrane-potential measurements in WT and mutant mtDNA strains; large dots denote biological-replicate means, and smaller dots represent single-cell values. **(B)** Maximum intensity projections of widefield microscopy images of cells containing indicated mtDNAs stained with 50 nM TMRM. Nuclear-encoded Su9-NG was used as a mitochondrial marker. Data in (A) are expressed as TMRM/Su9-NG intensity ratios, reflecting ΔΨ normalized to mitochondrial mass. WT cells treated with 10µM CCCP serve as a depolarized control. **(C)** Quantification of relative ATP levels in strains expressing mitochondrially targeted QUEEN-2m, obtained by determining fluorescence after excitation with 400 nm and 480 nm. WT cells in synthetic medium containing 2-deoxy-glucose instead of glucose provide an ATP-depleted control. **(D)** Representative images of QUEEN-2m–expressing cells, with pixel-wise 400 nm/480 nm ratio computed using the Image Calculator in FIJI. **(E)** Experimental outline of the mating of two haploid cells to generate a heterozygous zygote with a labelled P1 cell. **(F)** Representative widefield fluorescence images of zygotes expressing QUEEN-2m; parental cells share identical mtDNA in the upper row, or comprise WT and mutant mtDNA in the lower row. Bright-field panels indicate P1 and P2 cells. Scale bar for all images ((B), (D) and (F)) = 5µM. **(G, H)** Quantification of ΔΨ **(G)** and ATP levels **(H)** in zygotes, expressed as the ratio of values measured in the P2 cell relative to those in the P1 cell. Panels (A), (C), (G), and (H) were analysed for statistical significance by unpaired Student’s t-test on replicate means (*P *<*0.05; **P *<*0.005; ***P *<* 0.001).

To explore whether physiological differences between mitochondrial segments containing intact or mutant mtDNA persist locally within fused mitochondrial networks, we performed mating experiments to generate heteroplasmic zygotes and assessed local physiological differences (Fig 3E). To this end, we imaged QUEEN-2m ratios and TMRM intensities in zygotes formed by mating WT cells with cells containing either WT or mutant mtDNA. Importantly, mitochondria fuse in zygotes and matrix components equilibrate, while mtDNA and mtDNA encoded subunits show limited diffusion [10] (S3 FigA+B). To distinguish the mating partners in microscopy images, the cell walls of WT cells (P1) were labeled with concanavalin A (conA) linked to the far-red dye Alexa 633. Notably, in these mating assays, the P1 cell carried intact mtDNA but not the version encoding the Atp6–mNeonGreen fusion, thereby preventing misinterpretation of the signals originating from the fluorescent sensors. In zygotes formed by matings between cells both containing intact mtDNA, the fused mitochondrial network cells exhibited similar ATP levels and ΔΨ within the segments that were present within the P1 cell and the mating partner (P2) (Fig 3F–H and S3 FigC). In contrast, zygotes obtained from matings between cells containing intact mtDNA (P1) with cells containing mutant mtDNA (P2) retained marked differences. ATP levels and ΔΨ remained higher in the WT P1 cell compared to the mutant P2 parent cell. Although the mitochondrial network in the P2 parent cell exhibited increased ATP and ΔΨ relative to their haploid forms, likely reflecting some degree of metabolite exchange, mutant-derived compartments did not reach WT-like levels.

These observations support the idea that mitochondrial physiological parameters, such as ATP levels and ΔΨ, are locally influenced by the underlying mtDNA genotype and are not fully equilibrated within the fused network. This provides a functional basis for the sphere-of-influence model and suggests that local differences in physiology may enable the selective recognition of defective mitochondrial genomes.

### Functional Respiratory Complexes Are Required for Selective mtDNA Inheritance

We reasoned that if these local physiological differences underlie mtDNA quality control, then impairing the respiratory chain globally should abolish selection against mutant mtDNA.

According to the idea that ΔΨ or ATP levels generated by the OXPHOS complexes serve as signals to distinguish intact from mutant mtDNA, a general impairment of the electron transport chain in heteroplasmic cells should reduce and equalize ΔΨ and ATP levels across the entire mitochondrial network. This would obscure differences in mtDNA quality and thereby impair the selective detection of intact versus mutant genomes.

To test this hypothesis, we generated strains with nuclear deletions of subunits from either complex III or complex IV by deleting *RIP1* or *COX4*, respectively. We also deleted *ATP4* to examine the effects of impaired ATP synthase function. However, these *ATP4* deletion strains exhibited pronounced mtDNA instability, in line with previous reports [35], which precluded the analysis of mtDNA quality in this background. First, we examined ATP levels and ΔΨ of Δ*cox4* and Δ*rip1* strains containing WT mtDNA. As expected, ATP levels and ΔΨ in these strains dropped to levels observed also in strains containing mutant mtDNA (S3 FigE+F). Notably, following the mating of these cells with cells containing Δ*rip1* or Δ*cox4* and mtDNA^ic^, mtDNA^il^, mtDNA^Δ*cob*^, mtDNA^Δ*cox2*^ or mtDNA^Δ*atp6*^, we found that heterogeneities in the fused mitochondria of the resultant zygotes were no longer detectable (Fig 4A-D).

**Fig 4.**
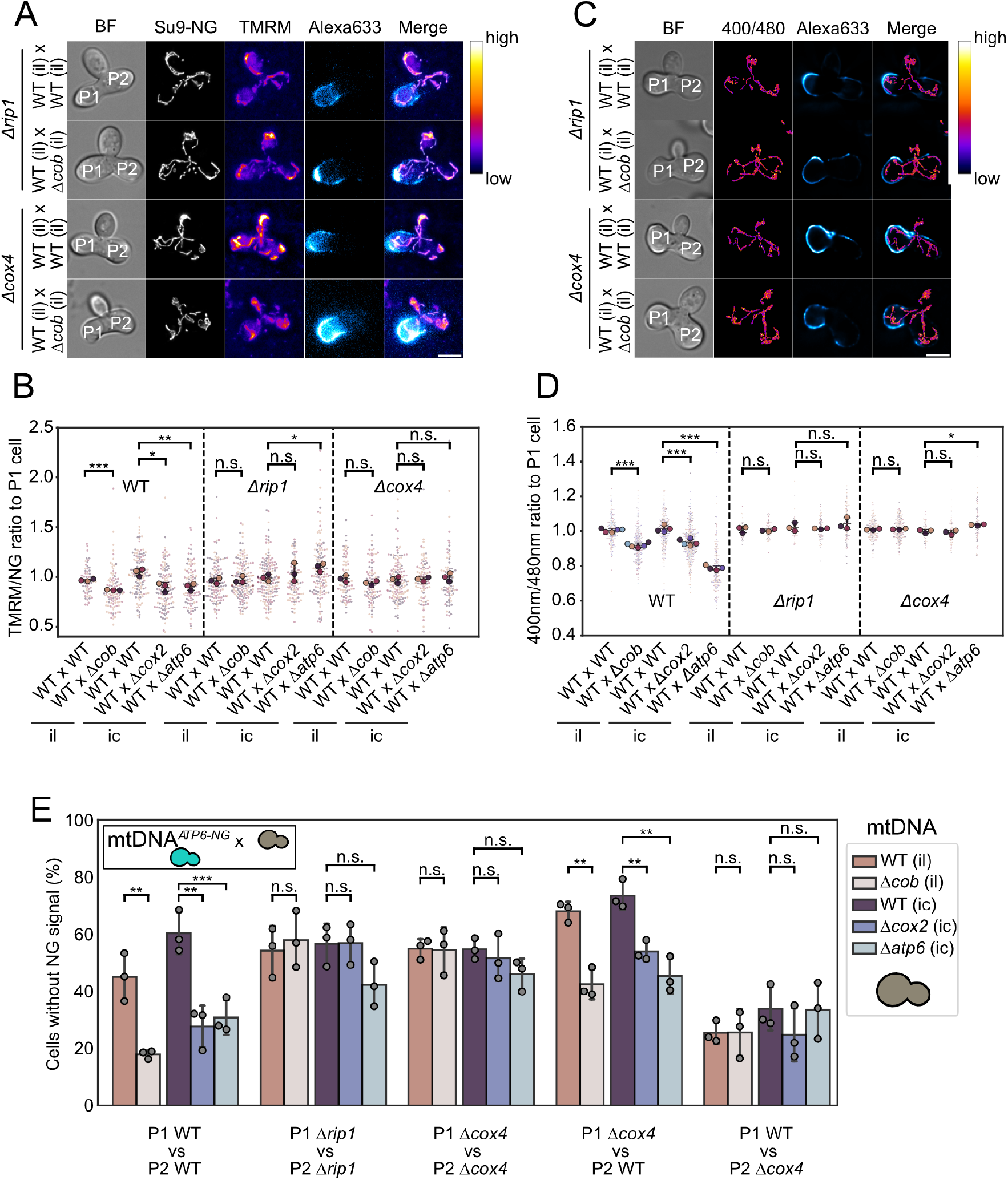
Functional Respiratory Complexes Are Required for Selective mtDNA Inheritance. **(A)** Maximum-intensity projections of widefield fluorescence images showing ΔΨ in heteroplasmic zygotes of Δ*rip1* and Δ*cox4* strains. **(B)** Superplot of P2 to P1 ΔΨ ratios in heteroplasmic Zygotes. **(C)** Maximum-intensity projections depicting ATP levels in heteroplasmic zygotes of Δ*rip1* and Δ*cox4* strains. **(D)** Superplot of P2 to P1 mitochondrial ATP level ratios in heteroplasmic zygotes. **(E)** FAST analysis results for homo- and heterozygous matings. Groups 1, 2 and 3 show zygotes with homozygous nuclear backgrounds (WT × WT and,Δ*rip1* × Δ*rip1* Δ*cox4* × Δ*cox4*; group 4 represents mtDNA^*ATP6-NG*^ Δ*cox4* × dark (*COX4* ^+^), and group 5 represents mtDNA^*ATP6-NG*^ (*COX4* ^+^) × dark Δ*cox4*. Panels (B), (D) and (E) were analysed for statistical significance by unpaired Student’s t-test on replicate means (*P *<*0.05; **P *<*0.005; ***P *<* 0.001).

We used flow cytometry to assess whether respiratory-deficient mutants continued to express Atp6-NG, a prerequisite for analysing mtDNA quality control during FAST. Fluorescece was observed in this cells, however, the proportion of non-fluorescent cells modestly increased in Δ*rip1* (7.4%) and Δ*cox4* (6.7%) strains compared to WT (2.0%) (S4 Fig A). We then performed FAST analyses to investigate the segregation of intact and mutant mtDNA in the progeny of heteroplasmic zygotes lacking *RIP1* or *COX4*. Notably, by using mtDNA^*ATP6-NG*^, the FAST assay enabled us to directly detect the presence of intact mtDNA even in respiratory-deficient Δ*rip1* or Δ*cox4* mutants, which is an advantage not afforded by our previous pedigree analyses, which relied on respiratory growth as a proxy for mtDNA integrity [10]. In the FAST mating experiments, purifying selection against Δ*cox2* or Δ*cob* mtDNA was abolished in Δ*rip1* and Δ*cox4* backgrounds, resulting in an approximately equal distribution of cells expressing green fluorescence and those lacking fluorescence (Fig 4E).

Notably, FAST analyses examining the competition between mtDNA^Δ*atp6*^ and mtDNA^*ATP6-NG*^ in Δ*rip1* and Δ*cox4* backgrounds revealed a lower proportion of non-fluorescent cells lacking mtDNA^*ATP6-NG*^. However, Δ*rip1* and Δ*cox4* strains harboring mtDNA^Δ*atp6*^ exhibited pronounced growth defects relative to the Δ*rip1* and Δ*cox4* strains carrying intact mtDNA^*ATP6-NG*^ (S4 FigB+C). This likely accounts for the underrepresentation of non-fluorescent cells in the mating experiments, as these cells are outcompeted by haploid progeny carrying the intact mtDNA^*ATP6-NG*^. The pronounced growth defect of Δ*cox4* and Δ*rip1* strains harboring mtDNA^Δ*atp6*^ likely stems from their inability to establish a ΔΨ-neither via electron transport chain activity nor through reverse operation of the ATP synthase.

To further investigate the role of respiratory function in mtDNA quality control, we performed a series of asymmetric mating experiments using the FAST assay, where either one of the mating partners harbored a deletion of the nuclear-encoded *COX4*. The P1 cell always contained mtDNA^*ATP6-NG*^, while the P2 cell carried either intact or mutant mtDNA (S4 FigD).

When the P1 cell contained the *COX4* deletion, we observed an increased proportion of non-fluorescent progeny (dark cells) in matings against both intact mtDNA^il^ and mtDNA^ic^. This suggests that, in the absence of respiratory activity in P1, intact mtDNA in the P2 cell is preferentially retained, even though the P1 cell also contains intact mtDNA (Fig 4E). In contrast, when both P1 and P2 cells were respiratory-deficient-either because both lacked *COX4* or because one carried a *COX4* deletion and the other harbored mtDNA deletions, mtDNA segregation was approximately 50:50, indicating a lack of selective pressure.

Conversely, when the *COX4* deletion was introduced into the P2 cell, we observed a strong bias toward fluorescent progeny in matings against P2 cells carrying intact mtDNA^il^ or mtDNA^ic^. This suggests that nuclear-encoded respiratory defects can promote selection against functional mitochondrial genomes in our mating assay. Notably, when P2 cells carried both a *COX4* deletion and mutant mtDNA (mtDNA^Δ*cob*^, mtDNA^Δ*cox2*^, or mtDNA^Δ*atp6*^), selection for the fluorescent P1 genome was still observed, but not more pronounced than in the absence of the *COX4* deletion. This is consistent with the notion that the loss of respiratory competence is already conferred by the mutant mtDNA, rendering the additional deletion of *COX4* functionally redundant in this context.

These results reinforce the model that mtDNA quality control relies on functional respiratory activity, and that both nuclear- and mitochondrial-encoded components of the electron transport chain are essential for selective mtDNA maintenance. Notably, in our mating assay, yeast zygotes formed from parental cells with or without a functional respiratory chain—due to the presence or absence of nuclear-encoded *COX4* —select against intact mtDNA variants when these reside in mitochondria rendered respiratory-deficient by a nuclear mutation. This highlights that mtDNA selection can act in a relative manner, requiring functional differences between mitochondrial populations to drive quality control.

## Discussion

In this study, we introduce FAST, a novel pipeline to analyse the segregation behaviour of mtDNA variants in heteroplasmic *S. cerevisiae* zygotes. Using this approach, we show that cells can distinguish between functional and mutant mtDNA genomes lacking components of complexes III, IV, or the ATP synthase, and preferentially maintain intact genomes in the resulting population. Within a fused mitochondrial network, regions containing mutant or WT mtDNA exhibit distinct physiological properties, such as differences in ATP levels and ΔΨ. Furthermore, we find that efficient mtDNA quality control requires nuclear-encoded components of the respiratory chain, highlighting that selection is tightly linked to respiratory function.

Interestingly, our segregation analyses combined with growth analyses and simulations reveal that intracellular selection against mtDNA^Δ*atp6*^ is weaker than against mtDNA^Δ*cob*^ or mtDNA^Δ*cox2*^. This finding is reminiscent of findings in mammalian systems, where a weaker selection against mutation in the *ATP6* gene was observed [9, 28]. We propose that this may be linked to the structural role of dimeric ATP synthase in shaping cristae morphology. Loss of *ATP6* disrupts dimer formation of the ATP synthase and compromises cristae organization, which may impair the spatial segregation of mitochondrial gene products, the so-called “sphere of influence”. Increased mixing of gene products between mitochondria harboring functional and mutant genomes could blur the physiological differences needed to drive selective maintenance of intact mtDNA.

Nonetheless, selection against mtDNA^Δ*atp6*^ is still detectable. Prior studies suggest that even in the absence of Atp6, partial ATP synthase assemblies and rudimentary cristae can persist [18], possibly allowing for a weakened but functional compartmentalization that supports some degree of mtDNA discrimination.

Our findings also reveal that mtDNA selection depends on a functional respiratory chain: in cells lacking nuclear-encoded components such as *COX4* or *RIP1*, selection against mutant mtDNA is abolished. Notably, we observe that even WT mtDNA can be counter-selected when residing in mitochondria with impaired respiratory function. In matings between two cells carrying WT mtDNA, one of which lacks *COX4*, the mtDNA from the respiratory-deficient parent is selectively lost. This suggests that local respiratory activity serves as a signal for mtDNA quality control, acting not only as a requirement but also as a determinant of selection direction.

These findings support the sphere of influence model in which mitochondrial function, likely through outputs such as ATP production or ΔΨ, drives local mtDNA selection. While the mitochondrial network is continuous, our data demonstrate that regions can maintain distinct physiological profiles depending on the quality of a nearby mtDNA copy. Indeed, we detect spatial differences in ΔΨ and ATP levels within fused zygotic mitochondria, and propose that similar heterogeneity exists in other heteroplasmic contexts. Although such compartmentalization seems at odds with mitochondrial fusion and diffusion, prior studies have reported similar local variation in ΔΨ [36–39], but without linking it to mtDNA content.

A major unresolved question is how mitochondrial physiology is translated into differential maintenance of mtDNA. One possibility is that local reductions in ATP or ΔΨ impair the import of nuclear-encoded proteins required for mtDNA replication or maintenance, thereby preventing propagation of defective genomes. Such a mechanism has been proposed in *Drosophila*, where local depolarization prevents import of replication factors [5, 7]. Beyond direct effects on import, reduced ΔΨ may also trigger signaling pathways, for example, via phosphorylation of the RNA-binding protein LARP by the kinase Pink1, which inhibit local cytosolic protein translation on mitochondria. However, this pathway is absent in *S. cerevisiae*, which lacks both Pink1 and LARP. This suggests that yeast cells may employ analogous but mechanistically distinct strategies to couple mitochondrial function to mtDNA quality control, likely involving yet unidentified factors.

Alternatively, selection may be enforced by differential transport of mitochondrial fragments during budding. The adaptor protein Mmr1, which links mitochondria to Myo2, has been implicated in directing functional mitochondria to daughter cells [40–42]. Spatial segregation of functional and non-functional mitochondria based on their physiological output could reinforce selective inheritance during cell division. However, the molecular mechanisms that allow Mmr1 to distinguish mitochondrial quality remain to be determined.

Another open question is whether selection mechanisms vary by mutation type. Our study focused on protein-coding gene deletions, but it remains to be tested whether similar selection occurs against mutations in mitochondrial tRNAs or rRNAs-lesions commonly found in human mitochondrial diseases. FAST now offers the opportunity to systematically explore how a broad range of mtDNA mutations affect inheritance dynamics in a high-throughput and quantitative manner.

Taken together, our results underscore the importance of mitochondrial function in guiding mtDNA selection, and establish *S. cerevisiae* as a powerful system to dissect the underlying mechanisms. The ability to model heteroplasmy, perform genetic manipulation, and apply scalable assays such as FAST opens the door to systematic dissection of mitochondrial quality control across diverse classes of mtDNA mutations.

## Materials and methods

### Yeast strains and Plasmids

All yeast strains are derived from the W303 background. Deletion of genes was performed using homologous recombination according to protocols described in [43]. Strains generated or used for this study can be found in Supplementary Table 1. The plasmids used are listed in Supplementary Table 2

### Yeast growth conditions

Yeast strains were kept on either rich media (1% yeast extract, 2% peptone, 0.04% adenine, 2% bacto-agar) with 3% glycerol (WT mtDNA/ functional respiratory chain) or synthetic media lacking arginine (0.67% yeast nitrogen base, 2% glucose, 0.192% drop-out mix minus arginine) (mtDNA^Δ*cob*^, mtDNA^Δ*cox2*^ and mtDNA^Δ*atp6*^) to ensure mtDNA maintenance prior to experiments. Unless stated otherwise, cells were grown overnight in rich media containing 2% glucose (YPD) at 30°C with 170 rpm shaking. In the morning, cultures were diluted to an *OD*_600_ of 0.1 and grown into early to mid log phase (*OD*_600_ 0.6-0.8).

### Fast pedigree analysis (FAST)

For mating, 500 µL of each strain in mid-log phase (*OD*_600_ 0.6–0.8) and of opposing mating type were mixed and transferred onto an YPD plate. Mating was conducted for 2 hours at 30°C.Following incubation, a tiny amount of cells was gently scraped from the plate using a pipet tip, resuspended in 30 µl of ddH_2_O, and pipetted onto a new YPD plate. The plate was tilted to allow the droplet to form a line, creating an optimal cell density for microdissection. The plate was then incubated at 30°C for an additional 30 minutes to promote sufficient growth of the first daughter cell from the zygote, allowing it to be detached from the zygote. After this period, 30 daughter cells were selected and relocated using a microdissection microscope (Singer Sporeplay+). Cells were allowed to grow into colonies for 20 h at 30°C. For flow cytometry, the formed colony was scraped off the plate with a pipette tip and resuspended in 300 µl sterile filtered SC medium. The samples were analysed with a BD Accuri C6 Plus Flow Cytometer with a limit of 10,000 events per biological replicate. Initial analysis and gating was performed using the floreada.ai tool followed by custom python scripts. To exclude false hits (e.g. dirt, agar) the events were gated for FSC-A (Forward-Scatter - Area) vs FSC-H (Forward-Scatter-Height).

### Cell growth determination

To determine cell growth, overnight cultures were diluted into fresh medium and incubated at 30 °C until reaching mid logarithmic phase (*OD*_600_ ~ 0.6-0.8). Cells were harvested by centrifugation, washed three times with sterile deionized water, and resuspended in YPD medium to an *OD*_600_ of 0.1. Cells were transferred into the wells of a 96-well plate, and growth was recorded at 30 °C on a SPECTROstar Nano (BMG Labtech) plate-reader at 20-minute intervals for up to 24 h, with continuous shaking at 800 rpm. Each strain was assayed in three technical replicates per plate. OD_600_ measurements were logarithmically transformed and fitted by linear regression to obtain the exponential growth rate (r, slope of the regression); doubling time (T_d_) was then calculated as

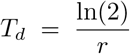

### Mating and stainings for microscopy

To distinguish P1 and P2 cells in zygotes during microscopy, WT cells (P1) were stained with conA-Alexa633 prior to mating. 1 mL of log phase cells (0.6 - 0.8) were spun down at 3000 g for 3 minutes. After decanting the supernatant, the cells were incubated at 30°C for 15 minutes in 500 µL 1x PBS containing 10 µg/mL conA-Alexa633 (ThermoFisher). Then, to mate the strains, 700 µL of each strain in mid-log phase were mixed in a reaction tube and centrifuged at 3000 g for 3 minutes. The supernatant was discarded and the cell pellet was resuspended in 50 µL of YPD medium. The suspension was applied to a YPD agar plate and gently spread into a line by tilting the plate. The liquid was then allowed to absorb/dry for 5–10 minutes. Plates were subsequently incubated at 30°C. After 90 minutes cells were scraped off the plate, and resuspended in 2 mL YPD. The *OD*_660_ was determined and 2 · 10^6^ cells were taken, centrifuged at 3000 g for 3 minutes and the supernatant discarded. The cells were then incubated at 30°C for 10 minutes in 1 mL 10 mM HEPES pH 6.7 supplemented with 50 nM Image-iT TMRM reagent (ThermoFisher), using a tube rotor (IKA Trayster digital). Then, the cells were washed twice in 1x PBS and loaded on a conA coated Ibidi slide (Ibidi GmbH) for imaging. Identical TMRM staining protocols were used for zygotes and single cells. For the DAPI staining of zygotes shown in S3 Fig A and B, conA-Alexa633 staining and mating were performed as described above. Subsequently, *OD*_660_ was determined and 2 · 10^6^ cells were collected by centrifugation at 3000 g for 3 minutes. The supernatant was discarded and the pellet was resuspended in 1 mL of SC medium supplemented with 2 µg/mL DAPI. Cells were incubated for 15 minutes at 30°C with gentle rotation using a tube rotor. After staining, cells were pelleted again, resuspended in 400 µL of SC medium, and transferred to a conA coated well of an Ibidi slide for imaging.

### Fluorescence Microscopy

Imaging was performed with a Nikon Ti2-Eclipse wide-field fluorescence microscope with a CFI Apochromat TIRF 100×/1.49 numerical aperture (NA) oil objective and a Photometrics Prime 95B 25mm camera. The microscope is encased by an environmental box to keep the temperature at 30°C. Yeast cells were immobilized on the surface of Ibidi 8-well µ-Slides (18-well slides were used for pedigree imaging) using pretreated wells with conA (1 mg/mL).

### Image processing and analysis

All microscopy images were deconvolved with the Huygens software (Scientific Volume Imaging). Analysis of the FAST experiments shown in S2 Fig A and B has been performed as described previously [12]. Budding cells and zygotes in microscopy images were automatically identified and segmented using YeastMate [44] within custom FIJI macros, followed by cropping of individual cells into single images. Mitochondrial segmentation was performed in three dimensions using Mitograph, and channel intensities were summed along the network coordinates provided by Mitograph (dimensions: xy= 0.11, z=0.2) [45]. Zygotes were subdivided into P1, P2, and daughter cells through the use of binary masks. These masks were generated semi-automatically with a custom macro using the YeastMate plugin in FIJI. Channel signals within the zygotes were then assigned to the corresponding parental or daughter cells based on the masks.

### Simulations

For the simulations in 2G, we adapted a previously established framework [12] and introduced a growth penalty for cells containing mutant mtDNA. First of all, we calculated the probability that, within a simulation step corresponding to the doubling time of the WT, a given cell divides and has a daughter cell *p*_*daughter*_ as follows:

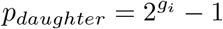

where *g*_*i*_ is the growth rate ratio (compared to WT) for the cell at time point *i*, which is calculated as follows:

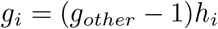

where *g*_*other*_ is the growth rate penalty of the strain crossed with WT (0.93 for mtDNA^Δ*atp6*^ and 1.0 for WT), while *h*_*i*_ is the mean of zeros and ones (representing the alleles of nucleoids) for the cell at time point *i*. Next, we ran the simulation 100 times, and for each run, we simulated 30 cells and calculated the average *h* for the colony. Note that all simulations were performed starting from a cell with 32 nucleoids distributed uniformly as sequence [1, 0, 1, 0, 1, 0…] (see S2 FigF). The other parameters of the simulation are *ngen, ndau*, and *nspl*. For the meaning of the parameters, we refer the reader to reference [12]. For 2G, we used the following values: *ngen* = 14, *ndau* = 11, and *nspl* = 5. Given the calculated WT strain doubling time of 1.48 hours, the number of generations *ngen* = 14 was chosen to simulate at least 20 hours, as in the experiments.

## Supporting information

**Supplementary Figure S1.**
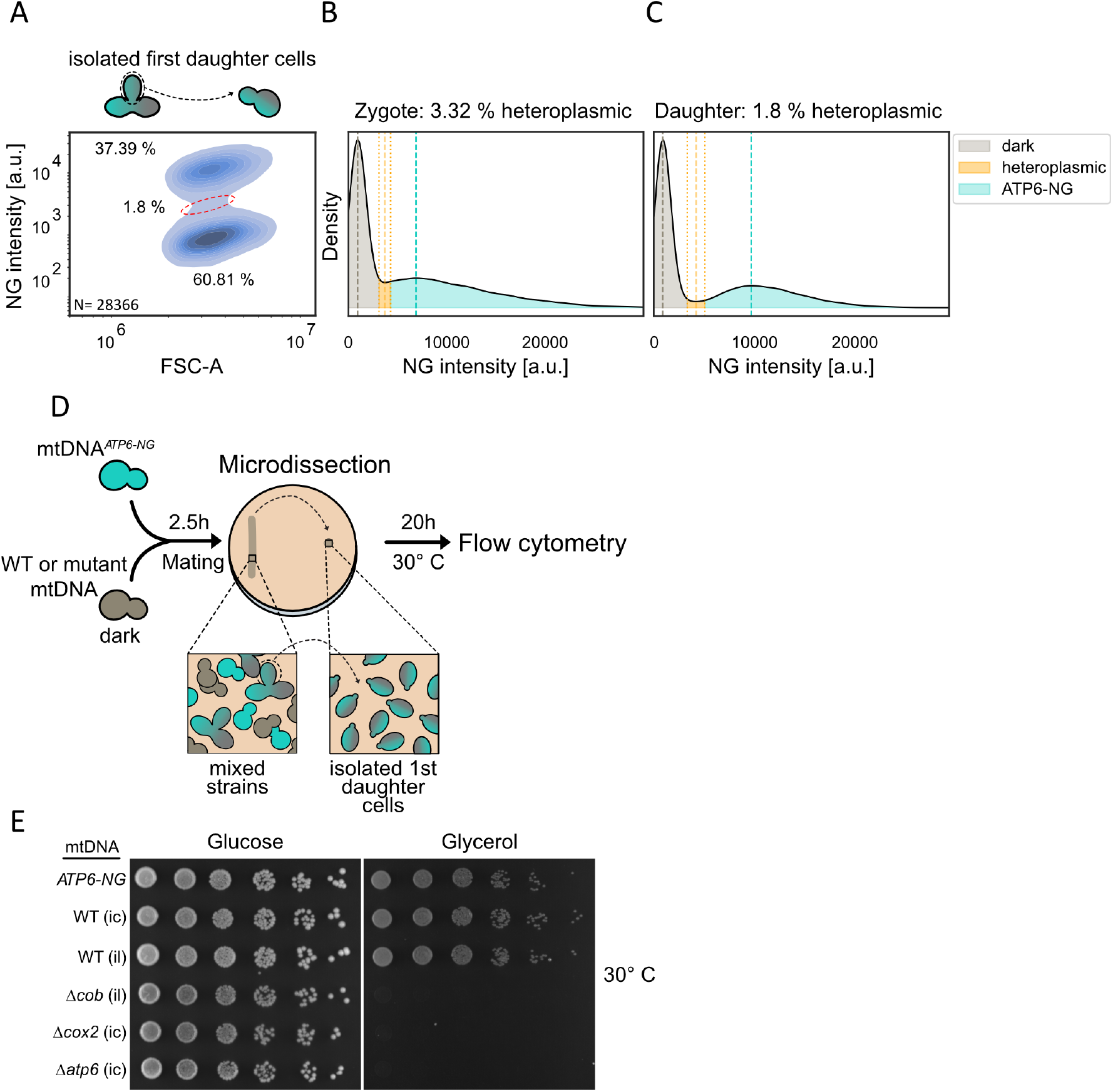
Proportion of heteroplasmic population in FAST using zygotes vs first daughters. **(A)** KDE plot of FAST analysis of mtDNA^*ATP6-NG*^ mated with mtDNA^ic^ when the first daughter cell of the Zygote was picked. Heteroplasmic cell population is highlighted by the red circle. Histogram of the NG intensity distribution among FAST population after 20 h of mtDNA^*ATP6-NG*^ mated with mtDNA^ic^ when **(B)** zygotes or **(C)** daughters of the zygotes were picked. Dotted lines mark the two peak maxima and the interpeak valley; shaded regions denote population assignments. The heteroplasmic window is defined symmetrically around the valley with a width of ±10 % of the interpeak distance. **(D)** Schematic of FAST experimental workflow. Schematic of the modified FAST workflow: instead of harvesting zygotes, first-division daughters are detached and relocated on agar. **(E)** Drop-dilution assay of mtDNA-variant strains was performed on fermentable (glucose) and non-fermentable (glycerol) rich media; plates were incubated at 30 °C and imaged after 48 h.

**Supplementary Figure S2.**
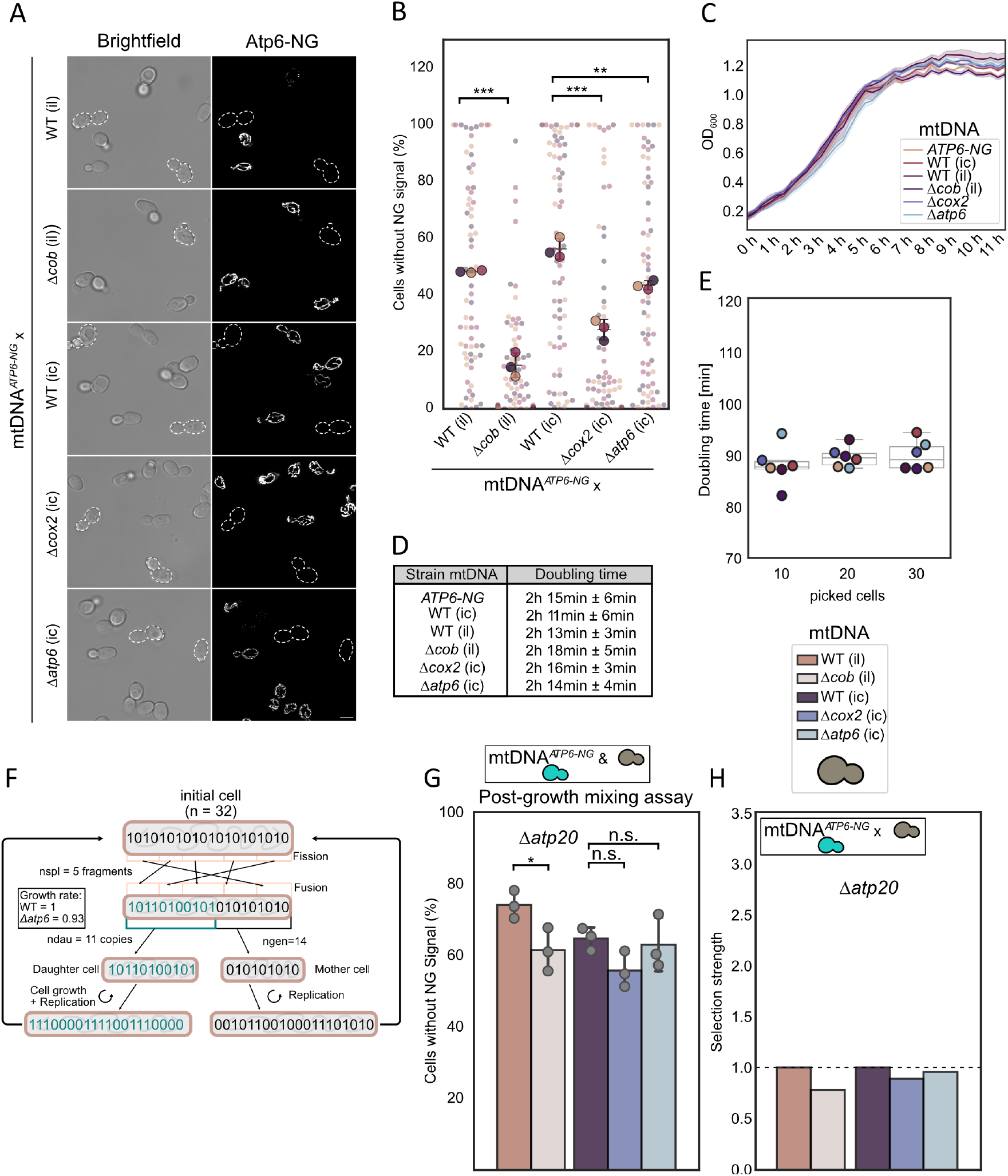
Confirmation of FAST by microscopy and determination of cell growth. **(A)** Representative widefield fluorescence microscopy images of microcolonies grown for 20 h following isolation of 30 first-division daughters from mtDNA^*ATP6-NG*^ *×* mtDNA^ic^, mtDNA^Δ*cob*^, mtDNA^il^, mtDNA^Δ*cox2*^, or mtDNA^Δ*atp6*^ matings. **(B)** Microscopy-based FAST quantification: individual daughter cells were arrayed on distinct plate positions, enabling each to form a microcolony over 20 h; colonies were then relocated to separate wells for imaging. Approximately 200 cells per well were scored for fluorescence status, with the ratio of fluorescent to non-fluorescent cells plotted as individual points in a swarmplot. The three big dots represent the mean of a single replicate. **(C)** Growth curves of all FAST strains: the solid curve represents the mean trajectory of three independent biological replicates, and the shaded region indicates the 95% confidence interval. **(D)** Mean doubling times for each strain, calculated from the growth curves in (C); errors represent one standard deviation (SD) across three biological replicates. **E** Doubling time of mtDNA^*ATP6-NG*^ cells grown on solid medium. Either 10, 20, or 30 individual cells were relocated to defined positions on the agar plate and cultured for 20 hours to allow microcolony formation. Subsequently, the resulting colonies were independently harvested into 100 µL of SC medium, and cell numbers were quantified by flow cytometry. Doubling times were calculated based on the known initial and final cell counts. Each data point represents a biological replicate. **F** Schematic of the workflow of the simulation used for Fig 2 G **(G)** Post-growth mixing assay of haploid Δ*atp20* cells. **(H)** Selection strength in Δ*atp20* cells calculated from FAST data (Fig 2 H) and post-growth mixing assay (S2 Fig D). Panels (B) and (G) were analysed for statistical significance using an unpaired student T-test was performed on the means of the replicates (*P *<*0.05; **P *<*0.005; ***P *<* 0.001).

**Supplementary Figure S3.**
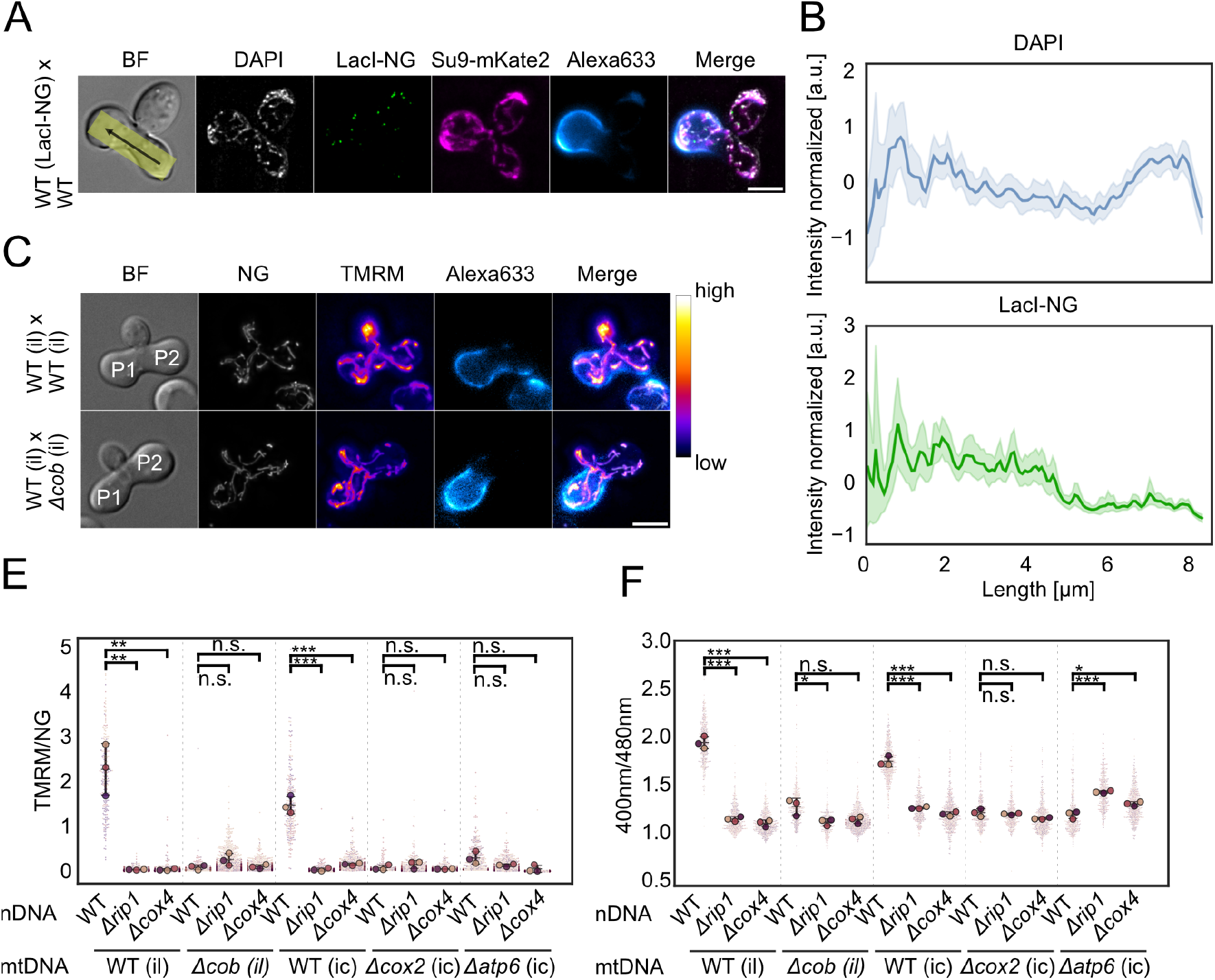
Phyisological parameters in cells lackign a functional respiratory chain. **(A)** Widefield fluorescence microscopy image of Zygotes. one parental cell expresses LacI-3xmNeongreen that can bind to the LacO repeats integrated in its mtDNA and mitochondrial targeted mKate2. Prior to mating its cell wall was stained with conA-Alexa633. The other cell does not express any fluorophores and its mtDNA lacks LacO repeats. Zygotes were stained with 2µg/ml DAPI to visualize the entire mtDNA pool of the cell. **(B)** Line profiles of DAPI and Laci-NG intensities of 50 Zygotes with a line width of 41 pixels as depicted in the BF panel of panel (A). Intensities were each normalized by z-score scaling **(C)** Representative widefield fluorescence micrographs of zygotes expressing Su9-NG (corresponding to Fig 3 G). Zygotes were stained with 50nM TMRM to assess ΔΨ. Parental cells contain either the same mitochondrial genome (upper row) or comprise WT and mutant mtDNA; P1 and P2 cells are indicated in the bright-field panel. Prior to mating, P1 cell walls were labeled with conA-Alexa633. Superplots of **(E)** mebrane potential levels or **(F)** ATP levels in zygotes of WT, Δ*rip1*, and Δ*cox4* nuclear background. The three big dots represent the mean of each replicate and small dots represent individual-cell values. Scale bar in (A) and (C) = 5µM. Panels (E) and (F) were analysed for statistical significance using an unpaired student T-test was performed on the means of the replicates (*P *<*0.05; **P *<*0.005; ***P *<* 0.001).

**Supplementary Figure S4.**
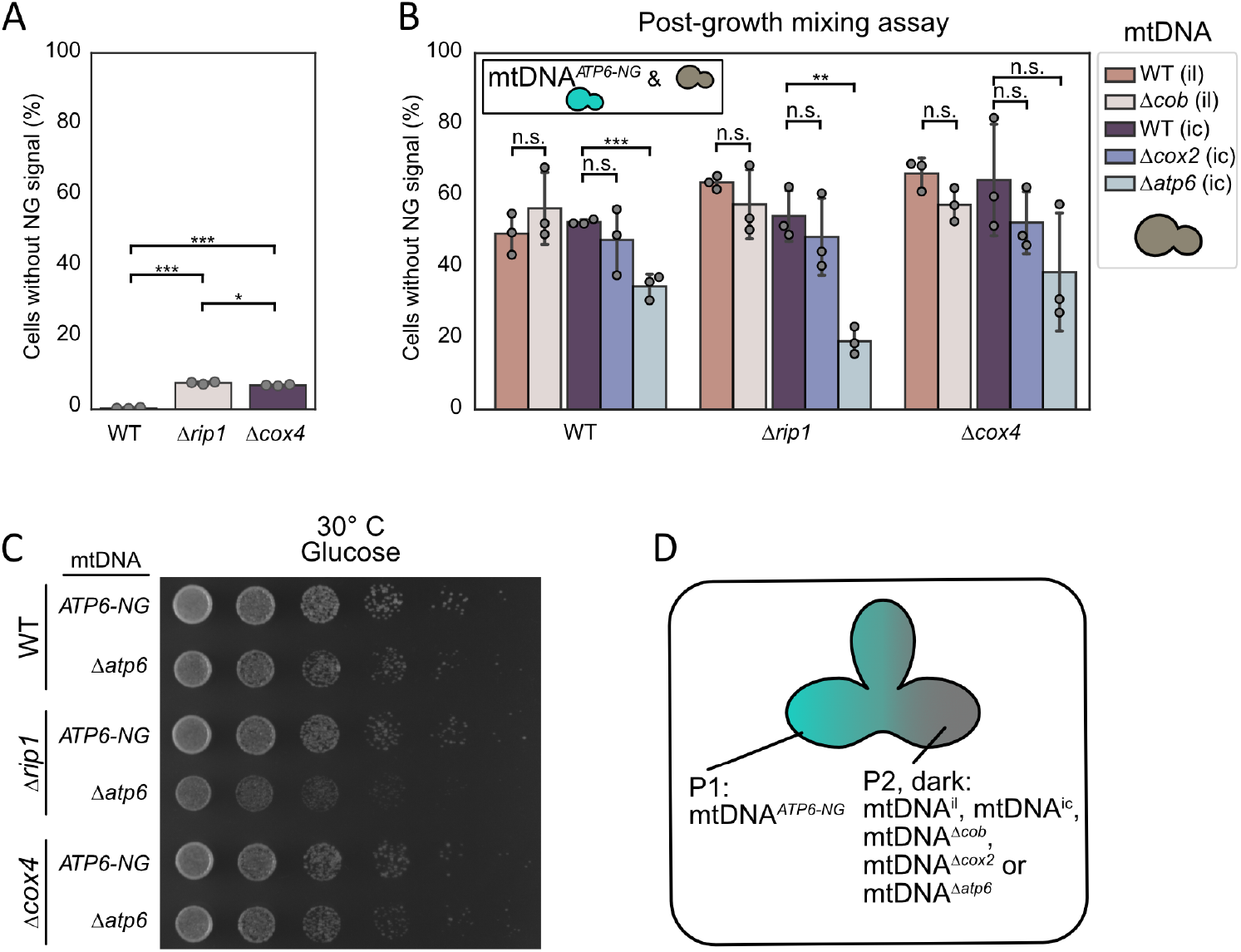
Growth phenotypes of mtDNA mutant strains with additional deletions in *RIP1* or *COX4*. **(A)** Fraction of cells within a mtDNA^*ATP6-NG*^ population without NG signal**(B)** Post-growth mixing assay of Δ*rip1* and Δ*cox4* nuclear-background strains compared to WT. **(C)** Drop dilution assay of mtDNA^*ATP6-NG*^ and mtDNA^Δ*atp6*^ strains in the WT, Δ*rip1* and Δ*cox4* nuclear background. Serial dilutions were spotted on YPD and incubated for 24 h at 30 °C. **(D)** FAST analysis results for homo- and heterozygous matings. Groups 1 and 2 show zygotes with homozygous nuclear backgrounds (WT × WT and Δ*cox4* × Δ*cox4*, data as in Fig 4 B); group 3 represents mtDNA^*ATP6-NG*^ Δ*cox4* × dark (COX4^+^), and group 4 represents mtDNA^*ATP6-NG*^ (COX4^+^) × dark Δ*cox4*. Panels (A) and (B) were analysed for statistical significance using an unpaired student T-test was performed on the means of the replicates (*P *<*0.05; **P *<*0.005; ***P *<* 0.001).

## Acknowledgments

We are grateful to AG Kunz and AG Robatzek for providing access to the flow cytometers. We thank Nadja Lebedeva for her technical assistance and to Verena Peterreins for cloning the QUEEN-2m plasmid. We also appreciate the valuable discussions and feedback from members of the Mokranjac lab and the Mito-Club. This work was supported by a DFG grant (OS 410/3-1) and a Human Frontier Science Program grant (RGP021/2023) awarded to C.O.. D.H. is supported by a DFG grant (HO 7333/1).

## Notes

### Competing Interest Statement

The authors have declared no competing interest.

